# Integrative Analysis of Phenomic, Genomic, and Transcriptomic to Identify Potential Functional Genes of Yaks in Plain and Plateau

**DOI:** 10.1101/2020.11.29.392167

**Authors:** Jiabo Wang, Jiuqiang Guan, Kangzhu Yixi, Tao Shu, Zhixin Chai, Jikun Wang, Hui Wang, Zhijuan Wu, Xin Cai, Jincheng Zhong, Xiaolin Luo

## Abstract

**Background:** The yak is an important source of livelihood for the people living in the Qinghai-Tibet Plateau. Most genetics detection studies have focused on the comparison between different tissues of different breeds, both living in the Plateau and in the plains. The genetic background and complex regulatory relationship have frequently puzzled researchers. In this study, we divided a population of 10 yaks into two subgroups, namely Plateau (living in the Plateau) and Plain (living in the plains). Phenomic, genomic, and transcriptomic analyses were used to reveal the molecular genetic type in the body weight, slaughter, and beef quality of yaks.

**Results:** We found a significant difference (P <0.01) between the third (60 days), fourth (90 days), fifth (120 days), and sixth (150 days) weights of Plateau and plain subpopulations. The difference in body weight was due to differences in kidney weight, meat weight, fur weight, and head weight. However, the beef quality was not significantly different. We identified 540 Differentially Expressed Genes (DEGs). Using weighted gene co-expression network analysis (WGCNA), we have constructed a co-express network, and the modules were strongly related to traits. In the genome-wide association studies (GWAS), we detected significant 156, 52, 33, 15, and 3 signals in the meat weight, head weight, fur weight, liver weight, and the last body weight traits. Based on the epigenome-wide association studies (eWAS) results, we created a link relationship between the DEGs expression level and genotype.

**Conclusion:** In summary, our study demonstrated the effectiveness and representative of multidimensional data from a finite number of yak populations. The study highlights the underlying way, as well as a related network, to yield new information on genome genetics and pathogen-host interactions of both the living Plateau and plain yak populations.

## Introduction

Yaks (*Bos grunniens*) are very important animals for the ecological environment and the people living in the Tibetan Plateau[1,2]. The Maiwa yak population is widely distributed in the Northeast of the Tibetan Plateau and ranks as the second major yak population in China[3–5]. Plateau adaptability (PA) has always been the major direction of yak genome research.

The approaches of mining genetics resources in yaks are differentially expressed genes (DEGs) detection and genome-wide association studies (GWAS) of quantitative trait loci (QTL)[6,7]. The DEGs or candidate QTL related to low atmospheric pressure, hypoxia, and high ultraviolet radiation in the Plateau were proven to play an important role in weight, growth, and meat quality[8–10]. However, the DEGs and candidate genes explained the insufficient genetics variance in the total phenotype variance. Such a biology problem, similar to that of other species named missing heritability, requires more effective methods and strategies to explain the interaction effect and other effects[11,12]. The network-assisted analysis, an emerging and promising method, improves functional DEG detection power and assists in creating new functional links between DEGs and complex traits for particular biological processes[13–15]. Creating correlated network approaches to cluster groups of genes and phenotype interactions will help us enhance our understanding of the pathophysiology of complex traits.

On the other hand, the multi-omics analysis offers a macroscopical interactive validation and mutual supplement to explain the molecular mechanism of production or adaptation traits[16–18]. There is a critical need for mining and classifying association links between such genes using multi-omics analysis. In this study, we sequenced the RNA expression data and re-sequenced the genomes data of 10 Maiwa yaks that either lived in the plains (Guanghan, 600 m altitude) or in the Plateau (Hongyuan, 3570 m altitude). Afterward, we integrated phenome analyses, QTL mapping (GWAS) by genome resequencing, and transcriptome profiling to filter and conclude the association link between genes that drive production traits in the yaks.

## Materials and methods

### Sample collection and phenotype measure

All 10 Maiwa yaks (*Bos grunniens*) had a similar health status and were approximately 3 years old with a similar weight (162.5±13.84 kg). The five randomly selected individuals were transported to Guanghan city (600 m altitude) and kept captive in pens, while the other individuals were kept in Hongyuan city (3570 m altitude) and pastured in the altitude steppe. Each individual was weighed with an empty stomach after the experiment started, on the 0th, 30th, 60th, 90th, 120^th^, and 150th day. On the 150th day, all yaks were slaughtered and measured with heart, liver, spleen, lung, kidney, bone, beef, fur, head, and foot weights. The three replicates in cooking loss rate, PH after 45 min, shear force, and the fresh color were used to compare the beef quality between these two groups. All clean tissue samples were divided into 1-2 cm^3^ sections and snap-frozen in liquid nitrogen for DNA resequencing and RNA sequencing (RNA-seq). Total RNA was extracted from muscle tissue using the TRIzol^®^ Plus RNA Purification Kit (Invitrogen, USA) in accordance with the manufacturer’s instructions. Ribosomal RNA (rRNA) was eliminated before sequencing using the Epicentre Ribo-zero™ rRNA Removal Kit in accordance with the manufacturer’s instructions.

### RNA-seq and co-expression network

After eliminating rRNA, cDNA sequencing libraries of total coding RNAs were generated using an mRNA-seq sample preparation kit (Illumina). Using the Illumina HiSeqTM 4000 platform, we obtained paired-end sequencing reads from 10 tissue samples. The sequencing reads were filtered using the fastp (version 0.19.8) software with default parameters and mapped to the reference genome (BosGru 3.0) using the hisat2 software (2.1.0)[19]. The different expression (DE) levels of the transcripts in each sample were measured using the StringTie-Ballgown analysis approach[20,21]. After removing the transcripts with low expression, in the DE analysis the fragments per kilobase of exon model per million reads mapped (FPKM) of transcripts were used to calculate the expression levels and the location was used as a covariation to compare the DE genes. |-log_10_(P value)|>2 and |log_2_(fold change)|>1 were used to filter DEGs. The R packages “clusterProfiler” and GO database “org.Bt.eg.db” were used to enrich DEGs relative to the biological process, cellular component, and molecular function[22]. Based on the expression levels of these DE transcripts, the functional modules were estimated using WGCNA[23]. The unsigned network was created to estimate the cluster group of DEGs.

### Whole-genome sequencing and SNP calling

Based on the same Illumina HiSeqTM 4000 sequencing platform, we obtained the fastq files with paired-end sequencing reads. The sequencing reads were filtered using the fastp software (version 0.19.8) with default parameters and mapped to the reference genome (BosGru 3.0) using the bwa software (version 0.7.17). The variations were identified using GATK (4.0.10)[24,25]. The total GVCF files of all samples were integrated into one VCF file, that was converted to a hapmap file using a perl script wrote by ourselves. The single-nucleotide polymorphisms (SNPs) were used to calculate population genetics and downstream association analysis. Markers with more than 10% missing rate were filtered. All remaining markers were imputed using the Beagle software (version 1.3.2)[26].

### GWAS and epigenome-wide association studies (eWAS) analysis

All GWAS and eWAS were performed in the genome association and prediction integrated tool (GAPIT) software with fixed and random model circulating probability unification (FarmCPU) model[27–29]. The top 3 principal components (PCs) and the indicator of environments were put into the model as fixed effects. The GWAS analysis tested five significant separate traits (meat weight, head weight, fur weight, liver weight, and last body weight). The eWAS analysis tested the expression levels of all DE genes as phenotype traits for detecting the association between DE genes and SNPs in the yak genome. In the GWAS and eWAS analyses, the false discover rate (FDR) <0.01 was used as the cutoff P value to filter the significant signals. The final circus plot was drawn using the RCircos package in R software[30].

## Results

### The separation of phenotype between Plateau and Plain

Following the separation of the two Plateau and Plain populations, the body weights of these two populations were significantly different (**Figure 1**). The red groups represent the yaks farmed at GuangHan (Plain). The blue groups represent the yaks farmed as Hongyuan (Plateau). The x axis is the time of weighing, at the 0th, 30th, 60th, 90th, 120th, and 150th day. At the first weight, there was no significant difference between Plateau and Plain (P value =0.208). However, as time increased, the difference gap became larger. With t-test, the P-values of the second, third, fourth, fifth, and sixth weighings are 0.026, 0.002, 6.43E-05, and 2.26E-05. We analysed the slaughter trait to determine what part of the yaks’ body caused this difference. Comparing slaughter traits between the Plain and Plateau groups (**Figure S1**), we found significant (P value<0.01) difference in kidney weight, meat weight, fur weight, and head weight. Although there was a significant difference between weights of yak bodies, there was no significant difference in the cooking percentage, PH after 45 min, shear force, and fresh color denoting meat quality (**Figure S2**).

**Figure 1.**
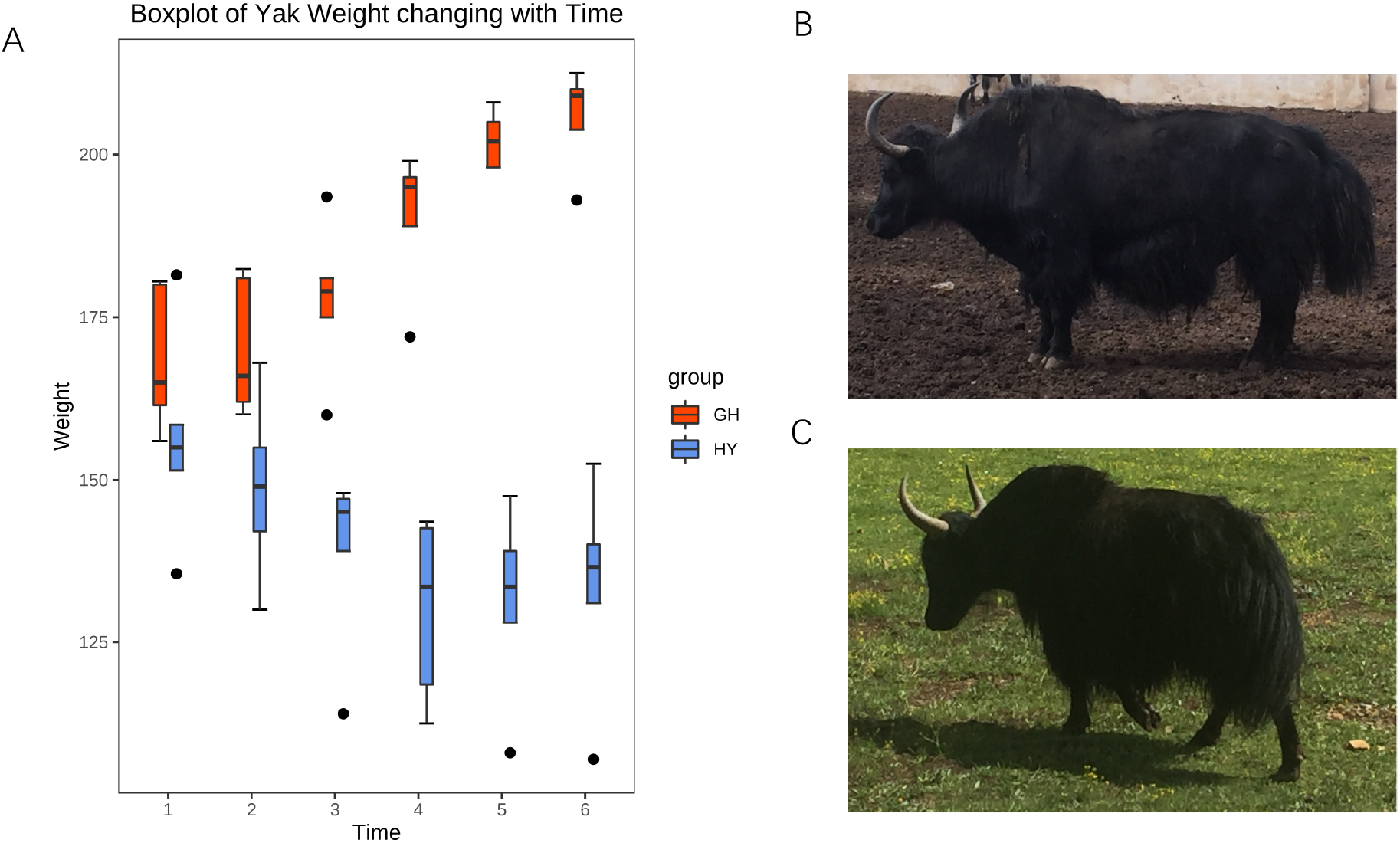
The Comparison of Yaks between Plateau and Plain. The red groups were the Yaks at GuangHan (Plain). The blue groups were the Yaks farmed as Hongyuan (Plateau). The x axis is the time of weighted at 0th, 30th, 60th, 90th, 120th and 150th days (A). The figure B was the Maiwa Yaks in the GuangHan (Plain) and the figure C was the Maiwa Yaks in the Hongyuan (Plateau).

### The transcriptome and genome sequencing

The sequencing transcriptome and genome data of Maiwa yaks in the Plateau (Hongyuan) and Plain (Guanghan) groups were obtained using the Illumina 4000 platform. After data quality control and filtering, we obtained 178G RNA-seq data and 292G resequencing data. The resequencing depth of each sample was more than 10X. All sequencing data were filtered using the fastp software, and the clean data were kept with Q30 read accuracy >91% (**Table S1**). The GC dinucleotide content was over 40% (DNA-seq) and 52% (RNA-seq). The reference genome (BosGru3.0) was individually mapped with DNA-seq and RNA-seq data. The mapping ratios (**Table S2**) were more than 97% (DNA-seq) and 88% (RNA-seq). After merging the GTF files of samples, a total of 86284 transcripts were assembled and saved as Ballgown format files for downstream analysis. After variance calling by GATK and filtering missing values >0.1, there were 872K SNPs kept as common genotypes of 10 Maiwa yaks. Afterward, we used Beagle to impute missing genotypes and filter the genotypes with minor allele frequency (MAF)<0.01.

### Transcriptome analysis

The FPKM expression levels of the transcripts were calculated based on the read counts using the Ballgown R package. Before DEG filtering, we removed some transcripts with a variance of <1 among the sample. A total 17841 transcripts were kept. The threshold -long_10_(P value)>2 and |log_2_(fold change)|>1 were used to identify DEGs between Plateau and Plain populations. A total of 540 DEGs were identified, 173 of these being upregulated and 367 downregulated (**Figure 2A)**. The known gene names (123) were signed in figure 2A and the unknown gene names (417) were marked as “.”. All DEGs were enriched into 1440 GO terms which included 1090 in biological processes (BP), 168 in cellular components (CC), and 182 in molecular function (MF). The top 15 significant GO terms (P <0.01) contained 94 DEGs (**Figure 2B**). The terms that enriched most DEGs (9) were regulations of the transferase activity, negative regulation of cell communication, and negative regulation of signaling. Those are the BP categories.

**Figure 2.**
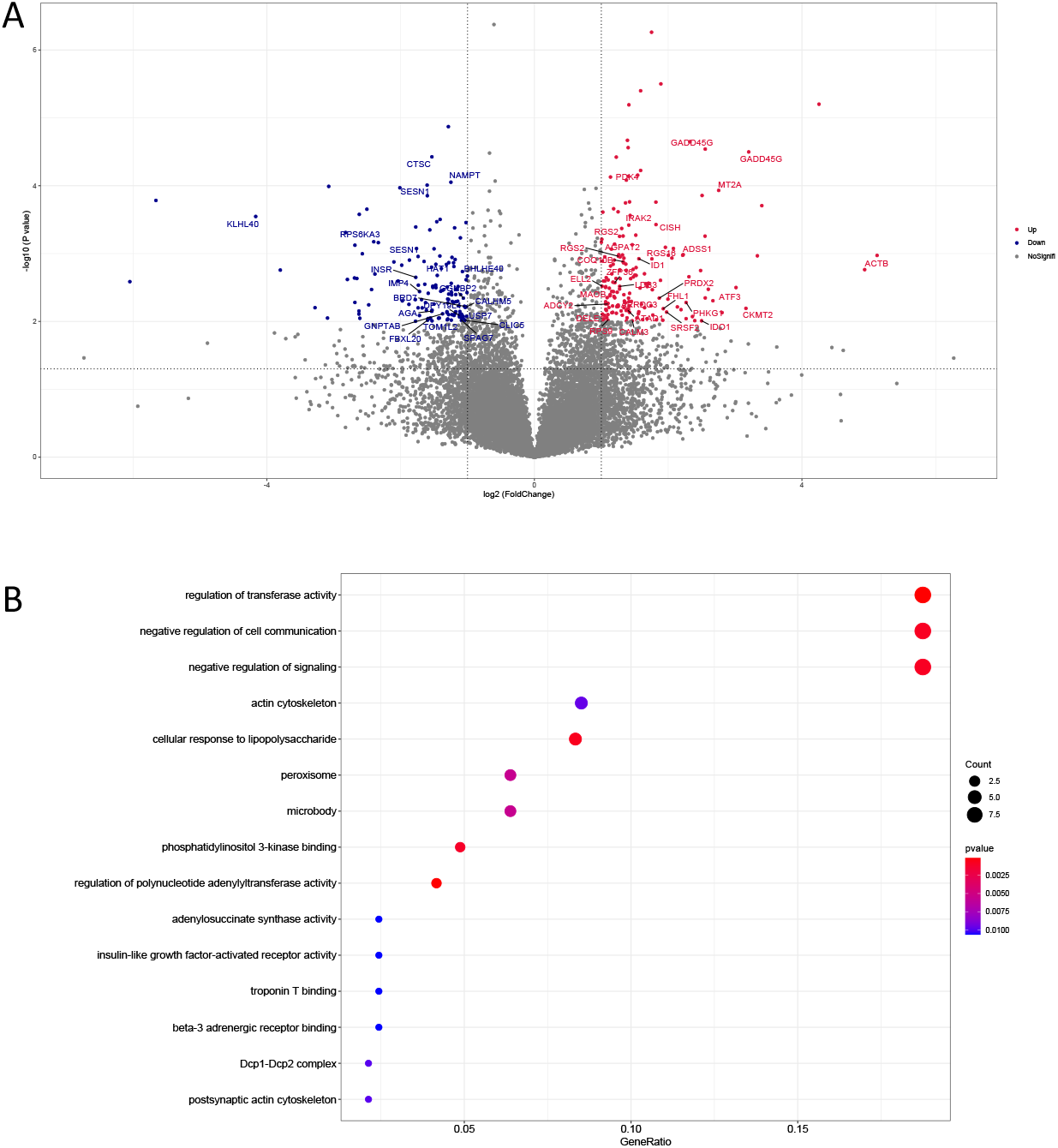
The Volcano Plot and GO analysis of Different Express Genes in Yak between Plateau and Plain. All transcripts were filtered by different express analysis. The red dots were up regulation DEGs, and the blue dots showed down regulation DEGs (A). -log_10_(P value)>2 and |log_2_(Fold change)|>1 were used to filtered DEGs. The figure B is the top 15 terms enriched DEGs in biological processes, cellular components and molecular functions. The size of dots showed the number of DEGs clustered in same terms. The color of dots indicated the P value.

Based on the expression levels of all DEGs between Plateau and Plain yaks, we used WGCNA to estimate an unsigned network for illuminating the functional modules of all DEGs (**Figure 3A**). The relationship between modules indicated the blue module close to the yellow module and the turquoise module was close to the brown module (**Figure 3B**). Although the unsigned network estimated four functional modules, they actually belong to two major functional large modules. We also compared the modules and traits (**Figure 3C**). The above value in the block is the correlation value and the value below is the P value. Dominant red shows a stronger correlation and dominant green shows a weaker correlation. Most of the weight traits had strong negative correlations to blue and yellow modules and strong positive correlations to turquoise and brown modules.

**Figure 3.**
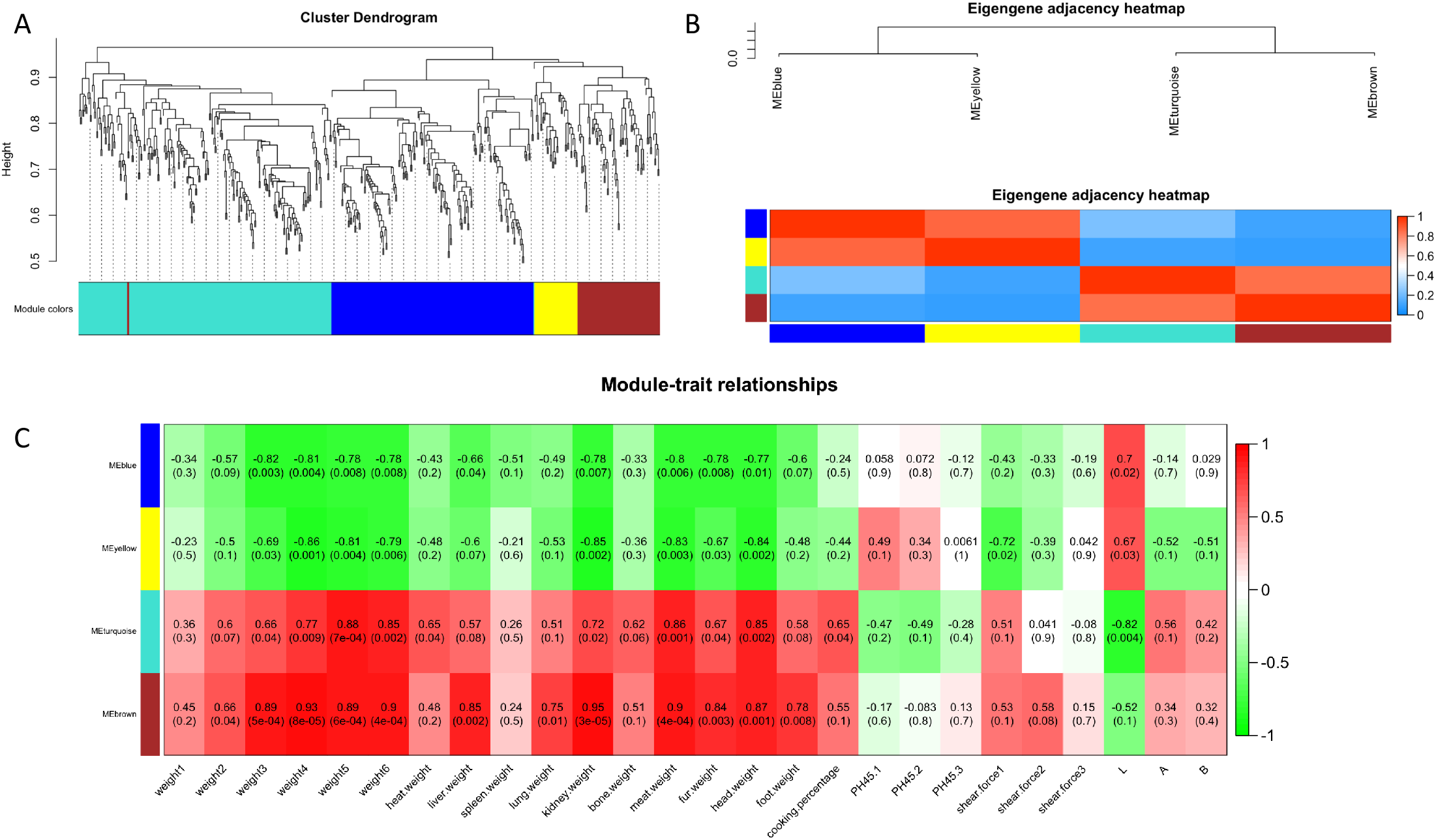
The WGCNA co-expressed network and the relationship between the modules and traits. In the A, all FPKM of DEGs were used to create the unsigned co-expressed network. The color shows the DEGs belonged to the module. Between each modules, the relationship indicated the blue module close to yellow module and the turquoise module close to the brown module (B). More red show stronger correlative and more blue color in the heatmap show weaker correlative. Between the modules and the traits, the heatmap indicated the relationship (C). The above value in the block is the correlation value and the below value is the P value. More red show the stronger correlative and more green show weaker correlative.

### Genome association detection

To integrate the analysis results and distribution from transcriptome profiling and QTL detection, we mapped the DEGs, QTL, and links of both DEGs and QTL onto the genome regions (**Figure 4**). The whole circle plot was composed of 8 circles. Using the FarmCPU method (**see Materials and methods**), we detected significant 156, 52, 33, 15, and 3 signals in the meat weight, head weight, fur weight, liver weight, and last body weight traits. The threshold was P <FDR(0.01). The FDR value was adjusted using the BH method. The common genes captured in the GWAS results and DEGs are shown in Table 1. In regard to meat weight, we filtered five common genes included *IDO1, RAD23B, DIXDC1, PSMD11*, and *DELE1* genes. In regard to head weight, we filtered three common genes included *IPO13, NOP14*, and *CTSC*. In regard to fur weight, we filtered two common genes included *STX12* and *RELCH*. To describe the link relationship between the DEGs and significant genes in the GWAS results, we used the expression levels of DEGs as the phenotype and genotype of the whole genome as variables in the eWAS model. All significant signals were considered to be significant link relationships (the centre circle of **Figure 4**).

**Figure 4.**
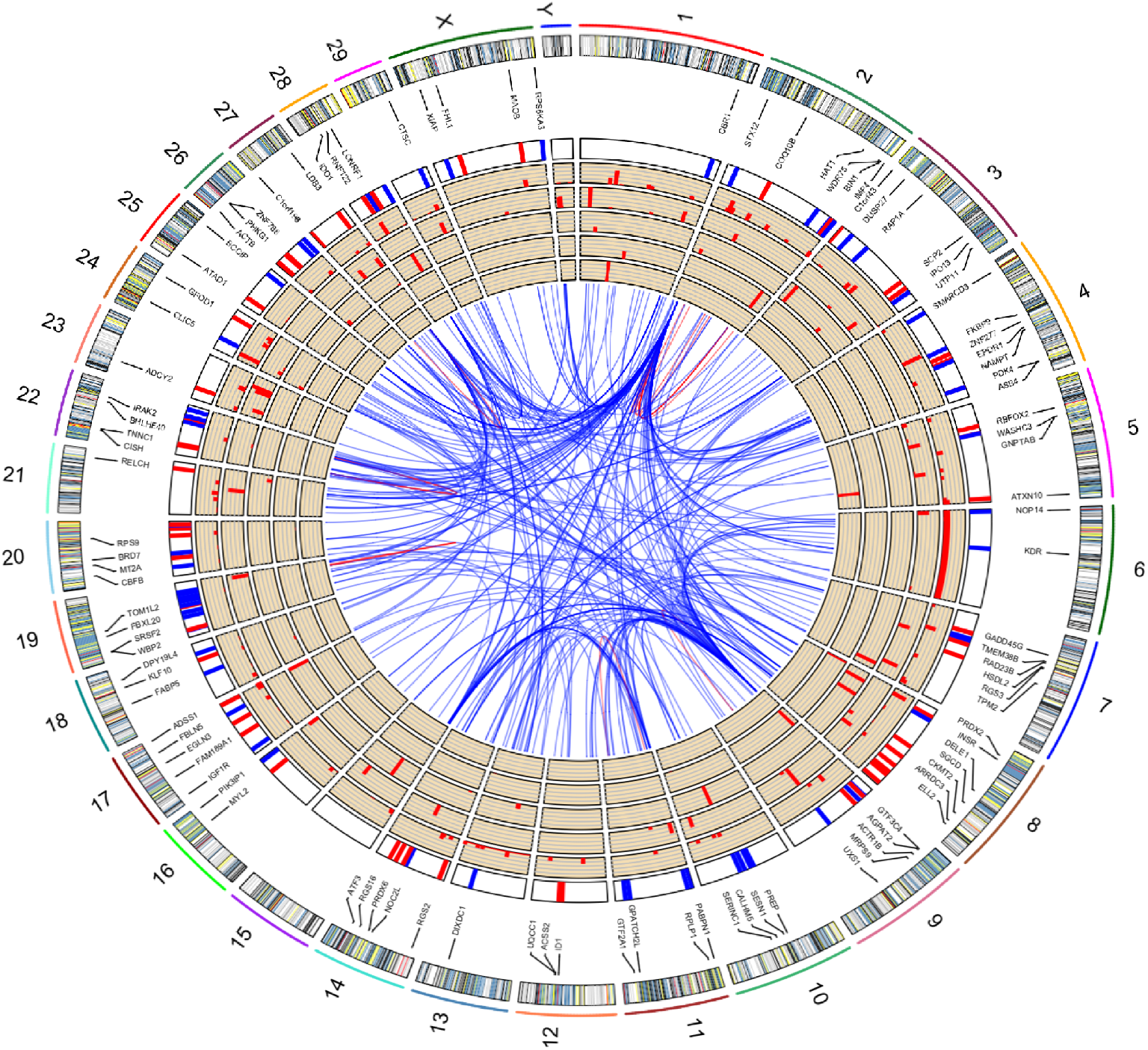
The distribution of mapped functional DEGs and significant association signals with traits in the Yak genome. The first outside loop is the distribution of functional DEGs in the yak genome. The color of bar show the DEGs clustered into modules. The second loop is the known gene name of DEGs. The third loop is the heatmap of expression levels of DEGs. The red showed the up regulation of DEGs in the Plain. The blue indicated the down regulation of DEGs in the Plain. The 4^th^, 5^th^, 6^th^ and 7^th^ circlos were significant GWAS results (the P value < FDR 0.01) of traits (Meat Weight, Head Weight, Fur Weight, Liver Weight and the last Body Weight). The inside link curves were the eWAS results between DEGs expression levels and genome genotypes. The blue curves showed the link relationship between the genes in different chromosomes. The red curves indicated the link relationship between the genes in same chromosome.

## Discussion

To explain the genetic adaptability, many studies have focused on the differences among the tissues of *Bos grunniens*, or the difference between the *Bos grunniens* and *Bos Taurus* lived in different altitude areas. However, regardless of the bias of the genetic background or complex environments, all caused immense noise for QTL mapping and DEG detection[17,31,32]. One doable approach to reduce these noises is to divide the yak population with similar genetic backgrounds and growing environments into two subpopulations respectively feeding in the Plateau and plain. In this study, we divide a Maiwa yak population with similar health, weight, and age into the Plateau and Plain groups.

Comparison of growing, slaughter, and meat quality traits between these two populations will help us to understand the subdivision of the different phenotype of individuals in the two groups with similar genetic backgrounds. In our phenome analysis, we found that after living longer, there are more differences in the body weights of yaks. From the slaughter traits analysis, the significant weight difference may be caused by kidney, meat, fur, and head weights. This indicates that the plain environment mainly increases the development of muscular and adipose tissue of the whole body of adult yaks. Comparison of meat quality traits, we did not find a significant difference (P<0.01). The altitude environments supply a few of the effects of changing the meant quality in adult yaks. While the cost of yak transported from Plateau to the plains is much cheaper than fattening yaks in the Plateau, breeding yaks in the plains is beneficial for improving yak bodies and meat weight, and for reduction of stress in the Plateau grassland.

In this study, we found 123 DEGs between Plateau and Plain yaks. Of these, the insulin-like growth factor 1 receptor (*IGF1R*) is the key gene in the progesterone-mediated oocyte maturation pathway[33]. As a receptor, IGF1R connects to IGF-1 and PL3K to improve oocyte maturation[34]. The *SGCD* and *MYL2* genes play important roles in the hypertrophic cardiomyopathy (HCM) pathway[35,36]. The RGS7 and IGF1R have also been reported to participate in the introgression of Mongolian yaks[37]. Most DEGs were enriched into three major terms in BP (regulation of transferase activity, negative regulation of cell communication, and negative regulation of signaling). The common genes between these three terms include several well-known candidate genes for Plateau adaptation and body development in humans and animals, such as the *ARRDC3, PRDX2*, and *IGF1R* genes[38–40]. The modules built with the WGCNA network were divided into two major modules (the genes in the blue and yellow modules are most downregulated expressed in the Plateau, while the genes in the turquoise and grown modules are most upregulated expressed in the Plateau). They are strongly related to yak weight traits (**Figure 3**). These upregulated expressed genes contained the *PIK3IP1, ELL2*, ARRDC3, *EPDR1, SGCD, PRDX2*, and *EGLN3*. The downregulate expressed genes contained the *IGF1R, RPLP1, EPDR1*, and *MYL2*. The co-expression network of these DEGs helped to understand the regulative relationship between each other.

Gene expressions in different locations or environments entail complex regulations and networks. The multi-omics information will help us understand these molecular mechanisms from a multidimensional perspective and classify the type of functional genes. In this study, the GWAS approach helped us reveal the associated markers of genome data with relative traits. The eWAS approach was used to build a network between these associated markers and DEGs of the transcriptome data (**Figure 4**).

## Supporting information

https://www.researchgate.net/publication/346061687_Supplymentary

https://www.researchgate.net/publication/346061794_Table_1?_sg=jUbp2K_Fj_2-SiDCgXFI7vD7KTgnn7ETFSCnjO5V2vZztF__yTkrtjxWrwbSDlUxM0C2xipU9wSHnn9V7gGl8oX

## Declarations

### Ethics approval and consent to participate

The protocol for Animal Care and Use, was approved by the Animal Ethics and Welfare Association of the Southwest Minzu University (No. 16053) and experiments were performed according to the regulations and guidelines established by this committee.

## Competing interests

The authors declare that they have no competing interests.

## Funding

This work is supported by the Program of Chinese National Beef Cattle and Yak Industrial Technology System (CARS-37) and Fundamental Research Funds for the Central Universities, China (Southwest Minzu University, Award #s 2020NQN26).

## Authors’ Contributions

JBW: Processed and analyzed sequencing data, implemented software, method and drafted the manuscript; JQG and TS: Participated in discussions regarding data analyses; JKW: Participated in discussions regarding interpretation of results; HW: Performed animal phenotyping and analyses for animal selection, collected samples; ZXC: Participated in discussions regarding interpretation of results; JCZ and XLL: Designed experiments, participated in discussions regarding data analyses. All authors made significant contributions editing the manuscript. All authors read and approved the final manuscript.

---

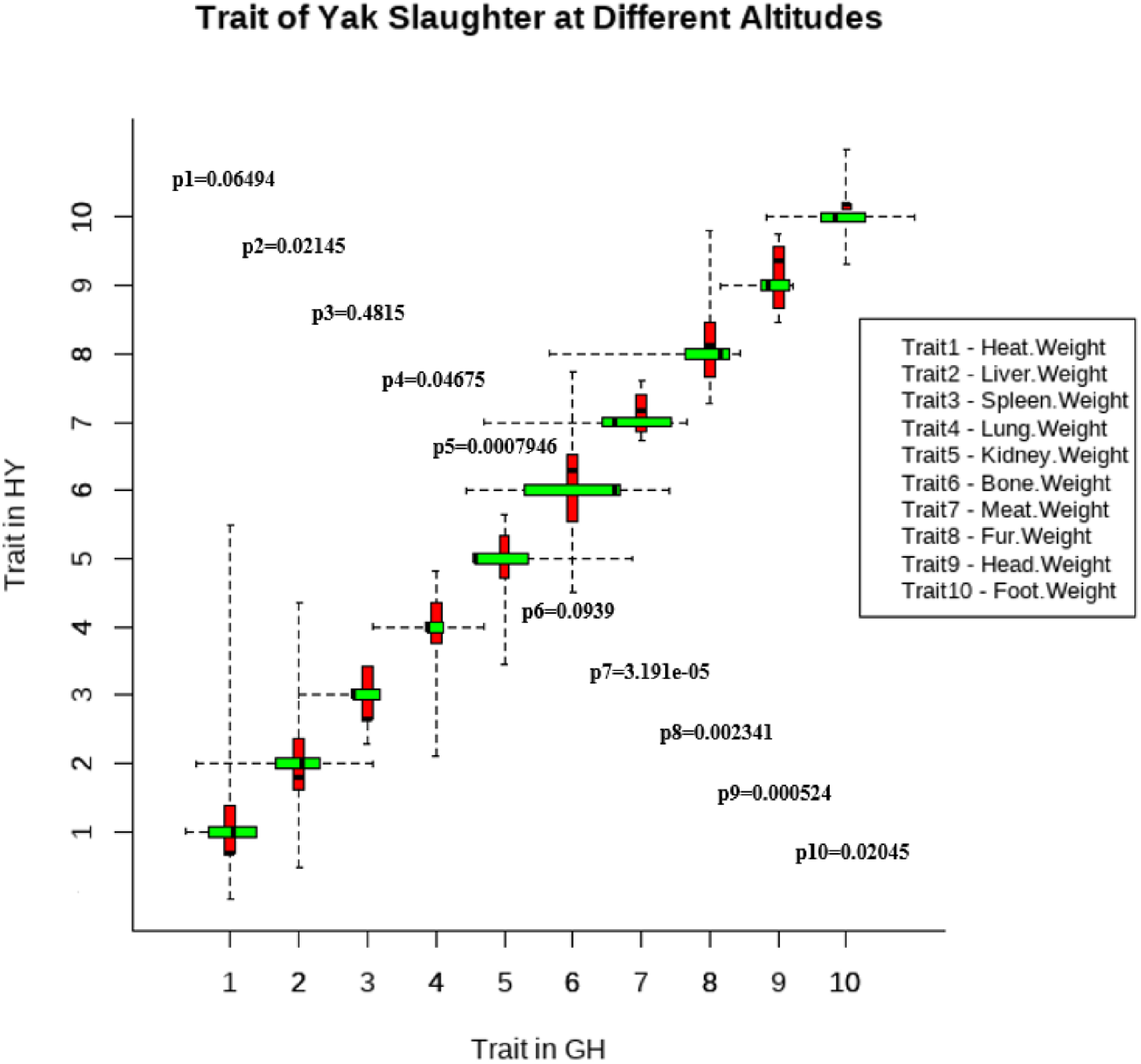

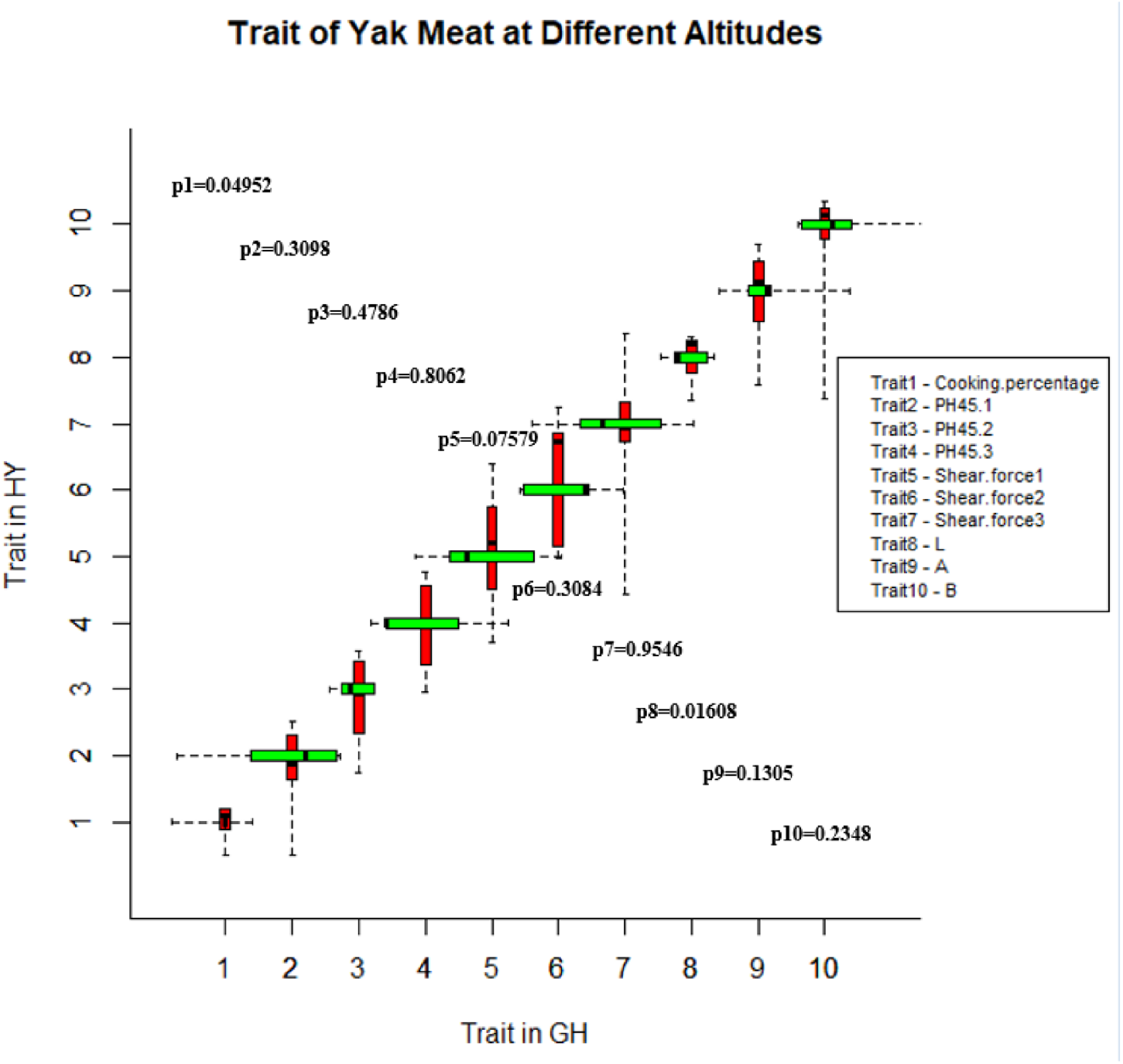

## Notes

### Competing Interest Statement

The authors have declared no competing interest.

