## Supplementary material for "Integrative Analysis of Phenomic, Genomic, and Transcriptomic to Identify Potential Functional Genes of Yaks in Plain and Plateau": https://www.researchgate.net/publication/346061687_Supplymentary

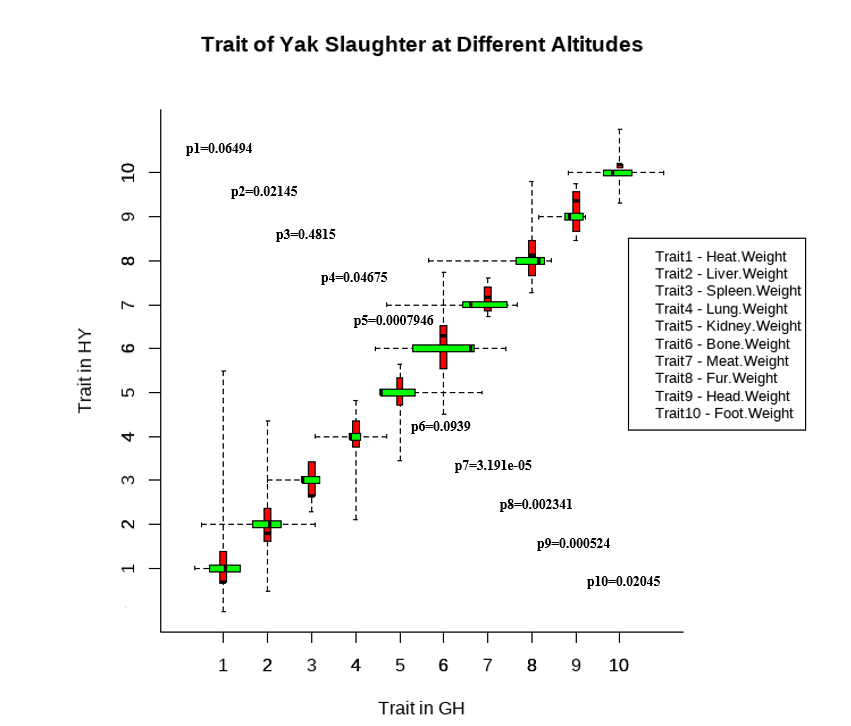

Figure S1. The comparison of phenotypes between HY (Plateau) and GH (Plain) in the Yak slaughter traits.

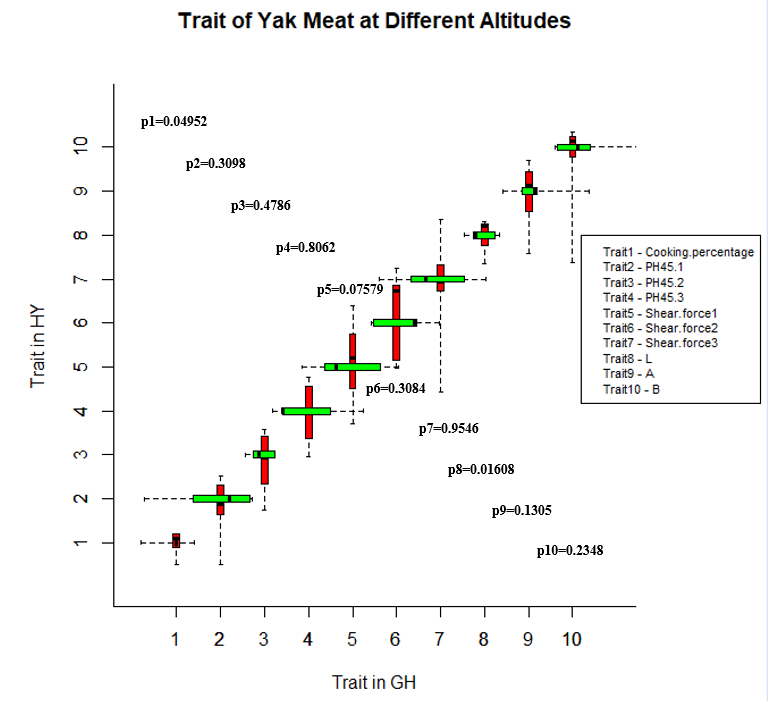

Figure S2. The comparison of phenotypes between HY (Plateau) and GH (Plain) in the Yak meat quality traits.

Table S1 The Sequencing quality of Maiwa transcriptome and genome

| Taxa | DNA Q30 | DNA Total bases | RNA Q30 | RNA Total bases |
| --- | --- | --- | --- | --- |
| M1 | 93.23% | 26.48G | 91.73% | 16.50G |
| M2 | 92.81% | 35.94G | 95.89% | 16.93G |
| M3 | 92.86% | 28.52G | 95.90% | 19.51G |
| M4 | 92.98% | 24.98G | 92.34% | 18.04G |
| M5 | 92.92% | 25.92G | 95.82% | 17.06G |
| M6 | 92.98% | 29.10G | 95.80% | 16.92G |
| M7 | 93.66% | 30.08G | 95.87% | 20.01G |
| M8 | 93.48% | 30.15G | 95.14% | 17.51G |
| M9 | 92.93% | 32.45G | 91.65% | 18.82G |
| M10 | 93.35% | 28.64G | 91.76% | 17.03G |

Table S2 The Mapping Raio of Maiwa transcriptome and genome

| Taxa | DNA Mapping | RNA Mapping |
| --- | --- | --- |
| M1 | 97.69% | 91.50% |
| M2 | 97.50% | 90.61% |
| M3 | 97.67% | 88.00% |
| M4 | 97.64% | 92.15% |
| M5 | 97.74% | 92.12% |
| M6 | 97.68% | 91.53% |
| M7 | 97.69% | 95.87% |
| M8 | 97.72% | 95.14% |
| M9 | 97.40% | 91.65% |
| M10 | 97.56% | 91.76% |
