## Supplementary material for "Integrative Analysis of Phenomic, Genomic, and Transcriptomic to Identify Potential Functional Genes of Yaks in Plain and Plateau": https://www.researchgate.net/publication/346061794_Table_1?_sg=jUbp2K_Fj_2-SiDCgXFI7vD7KTgnn7ETFSCnjO5V2vZztF__yTkrtjxWrwbSDlUxM0C2xipU9wSHnn9V7gGl8oX

| Numbers | Traits | Gene Name |
| --- | --- | --- |
| 5 | Meat Weight | IDO1, RAD23B, DIXDC1, PSMD11, DELE1 |
| 3 | Head Weight | IPO13, NOP14, CTSC |
| 2 | Fur Weight | STX12, RELCH |
| 0 | Liver Weight | - |
| 0 | Last Body Weight | - |

**Table 1.** The common genes between GWAS significant results and DEGs in 5 traits. The window size of significant SNP was 500kb. The common genes were filter with comparison of the window size of significant SNP and DEGs region.
